# KyDab – a comprehensive database of antibody discovery selection campaigns

**DOI:** 10.64898/2026.03.25.713450

**Authors:** Qingqing Zhou, Dawid Chomicz, David Melvin, Mark Griffiths, Sabrina Yahiya, Stephen T. Reece, Marguerite-Marie Le Pannerer, Konrad Krawczyk

**Affiliations:** Kymab, a Sanofi Company, Babraham Research Campus, Cambridge, United Kingdom; R&D, NaturalAntibody, Szczecin, Poland

## Abstract

Preclinical antibody discovery relies on progressive screening and down-selection of candidate antibodies from large immune repertoires, yet this critical process is poorly represented in existing public databases. Here we introduce KyDab (Kymouse Antibody Database), a well-curated database of antibody discovery selection data generated using standardized workflows on the Kymouse humanized mouse platform. The current release includes 11 Kymouse platform mice immunisation studies covering 51 immunogens, more than 120,000 paired heavy–light chain sequences, and binding measurements for a selected subset of experimentally characterized clones. By capturing full-funnel selection data with consistent metadata and both positive and negative experimental outcomes, KyDab provides a valuable data resource for the development and evaluation of artificial intelligence models for antibody discovery. KyDab is accessible at https://kydab.naturalantibody.com/, and the database will be continuously updated as new datasets become available.

## Introduction

Monoclonal antibodies have emerged as the fastest-growing class of biopharmaceuticals, with over 150 antibodies approved by the FDA to date^1^. Due to their superior specificity, potency, and safety profiles, antibody-based therapeutics are widely regarded as leading candidates for treating cancer, immune-mediated disorders, and infectious diseases^2^. In recent years, advances in artificial intelligence have begun to reshape antibody drug discovery workflows. Approaches such as virtual screening, affinity maturation, and generative design are increasingly integrated into early discovery, offering opportunities to accelerate development timelines and reduce costs^3^. These methods, however, rely heavily on large, high-quality datasets with experimentally validated outcomes to enable effective model training and benchmarking^4^.

To support the growing data needs of AI development, several public resources curate large antibody sequence datasets (e.g., Observed Antibody Space;^5^) and structural annotations (e.g., SAbDab;^6^). While these databases provide broad coverage, they do not capture the real-world process of therapeutic antibody discovery as practiced in an industrial setting. Modern discovery pipelines, including phage display, yeast display and animal immunisation, typically generate large pools of clones, followed by multiple stages of down-selection and affinity characterisation using assays such as surface plasmon resonance (SPR). Existing specialised therapeutic antibody databases, such as Thera-SAbDab^7^ and IMGT/mAb-KG^8^, focus primarily on the relatively small set of mature, clinically advanced antibodies. To our knowledge, no public resource currently provides full-funnel datasets that reflect the practical selection and screening steps used in real-world antibody discovery workflows.

Beyond these limitations in dataset representation, another critical gap concerns the scarcity and bias of affinity-characterized antibodies^9^. This is largely due to several factors - manual curation is labor-intensive, affinity values are often presented only as figures or partial tables, and negative screening data are rarely reported. Even in newer structural resources such as SAAINT-DB, which reviewed nearly 10,000 PDB entries, only ~1,400 non-redundant antibody–antigen affinities were identified^10^. Broader protein interaction databases, including SKEMPI v2.0^11^, AbBind^12^, Affinity Benchmark v5.5^13^ and PDBbind v2020^14^, catalog tens of thousands of measurements, but fewer than 2,000 involve antibody–antigen complexes. Moreover, because many databases draw on overlapping sources, the same affinity measurements are often repeatedly re-curated across databases rather than newly contributed. This scarcity of affinity-resolved antibody data remains a major bottleneck for AI-driven discovery.

To begin addressing these challenges, we present KyDab (Kymouse Antibody Database), a comprehensive database of antibody discovery selection datasets. KyDab provides systematically curated antibody sequences with comprehensive metadata and, where available, experimentally measured binding data. All datasets originate from Kymouse^15^, a state-of-the-art humanized mouse platform used for therapeutic antibody discovery. Consistent single-cell processing and standardized data analysis pipelines are applied to minimise technical variability and ensure comparability across campaigns. The current release spans 11 immunisation studies, comprising 51 unique immunogens, and 123,527 productive heavy–light chain pairs, with 1,657 clones associated with binding data. By making such campaign data available, KyDab establishes a resource that acts as a real-world benchmark for general-purpose antibody predictors. As such we expect that the database will foster development of more robust algorithms that can be applied in production in therapeutic programs.

## Materials and Methods

### Antibody Sequence Data

All antibody sequences deposited in this portal were generated from the Kymouse platform^15 16^, a well-established transgenic mouse system for producing antibodies with fully humanized variable regions. Each dataset is accompanied by a dedicated landing page accessible from the KyDab portal homepage, providing flexible download options as well as detailed information on immunisation, cell sorting, and sequencing protocols for each study. Sequences were initially processed and analyzed internally with an in-house bioinformatics pipeline, and subsequently re-annotated using RIOT^17^ prior to deposition in the portal to ensure consistent annotation across datasets and to avoid the release of potentially commercially sensitive information.

### Assay Data

Where available, following sequence analysis, a subset of clones were selected for expression and affinity testing. The specific assays employed in each study are described in the corresponding publications cited on the individual landing pages.

### Datasets collection

Summary of datasets are provided in **Table 1**.

**Table 1.** Datasets available in the KyDab portal.

| Studies | Immunogens | Binding Assay | Number of antibodies with binding measurements | Total Paired Antibody Sequences |
| --- | --- | --- | --- | --- |
| Pertussis | Pertussis toxin WT; Pertussis Toxin R9K E129A mutant | SPR | 422 | 11290 |
| Malaria CSP #1 | CSP-based immunogens | SPR | 124<sup>a</sup> | 18146 |
| Malaria CSP #2 | CSP-based nanoparticle immunogens | SPR | 110<sup>a</sup> | 22427 |
| Malaria CSP #3 | CSP-based nanoparticle immunogens | NA | 0 | 6401 |
| SARS-CoV-2 #1 | Sarbecovirus/beta coronavirus-based nanoparticles and soluble trimers | NA | 0 | 26379 |
| SARS-CoV-2 #2 | SARS-CoV-2 spike nanoparticles/trimer, RBD nanoparticle | HTRF | 249 | 9887 |
| Acinetobacter baumannii #1 | OMV cocktail | ELISA | 256 | 7383 |
| RSV #1 | Pre-fusion hMPV/RSV F protein-based nanoparticles, trimers and cocktails | NA | 0 | 10751 |
| RSV #2 | Stabilised Pre-F protein, Post-F protein, live/inactivated RSV | NA | 0 | 1099 |
| Influenza #1 | Stem-only HA immunogens and vaccine control | NA | 0 | 4106 |
| Typhoid #1 | Typhoid toxin E133A H160Q mutant | SPR | 496 | 5658 |

<sup>a</sup>Binding measurement data were generated by Antibody Dynamics.

### Availability

The datasets are available for researchers through our portal :

https://kydab.naturalantibody.com/

## Results

### Overview of the database

All datasets deposited in the portal are generated from mice derived from the Kymouse platform, a transgenic mouse platform developed by Kymab for discovery of therapeutic human monoclonal antibodies. These mice are genetically engineered to produce antibodies with variable heavy and light chain regions (VH and VL) of entirely human origin. Over the past decade, this humanized mouse platform has proven to be robust, stable, and versatile for therapeutic antibody discovery, with several antibodies derived from this platform having advanced to clinical trials^18 19^.

KyDab comprises 11 human antibody studies encompassing 51 unique immunogens generated from the Kymouse platform, targeting a diverse array of viral, bacterial, and parasitic antigens^15 16^. In general, immunogens were either selected from well-established commercial antigen products or designed by structural biologists with expertise in pathogen-specific immunogenic structures. Following immunisation with the respective immunogens, sera were collected and assessed for reactivity using screening assays such as ELISA to evaluate immune responses. Mice demonstrating positive immune responses were selected for further processing.

From each selected mouse, spleen, lymph node, and/or bone marrow cells were harvested and subjected to flow cytometry-based single-cell sorting to isolate antigen-specific B cells for downstream sequencing. The resulting paired VH-VL sequences underwent analysis and down-selection through an in-house bioinformatics pipeline that is similar to the Immcantation framework^20^. Sequences were clustered into lineages based on inferred V and J gene usage and CDR3 sequence identity, with representative clones from the more expanded lineages preferentially selected. Additional selection criteria included somatic hypermutation frequency, evidence of sequence convergence, developability assessments, and overall sequence diversity. The scale of clone selection varied across studies according to specific research objectives. In most studies, selected clones were recombinantly expressed and further characterized through in vitro affinity assays to assess binding to their corresponding antigens, as illustrated in **Figure 1**. The portal provides comprehensive data including all paired antibody sequences, germline gene annotations, complete metadata (mouse ID, tissue source, immunogen identity), and where available, affinity measurements and antigen sequences in FASTA format. To ensure uniform representation of the data, we post-processed raw read files using RIOT^17^, a sequence annotation tool that translates nucleotide sequences into amino acids and aligns them to the IMGT scheme.

**Figure 1.**
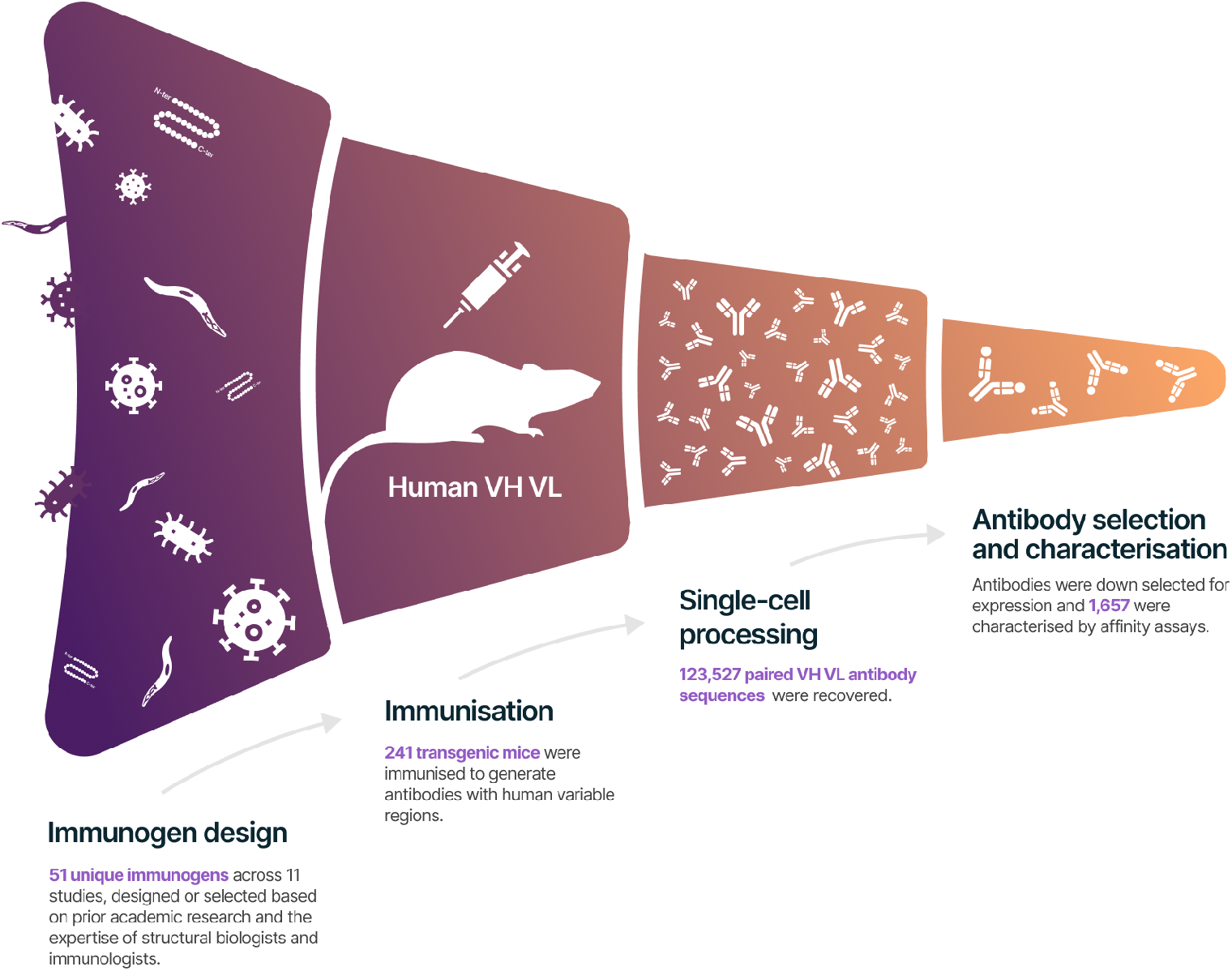
Overview of the KyDab data generation pipeline. All antibodies in KyDab were generated from the Kymouse humanized immune mouse platform. Following immunisation, antigen-responsive mice were selected and lymphoid tissues were harvested for single-cell sorting and sequencing. Resulting sequences were analyzed and down-selected to prioritise high-potential clones, which were often recombinantly expressed and further characterised by in vitro affinity assays. Both positive and negative binding data are made available on KyDab.

For friendly user experience, KyDab also features simple structured summaries, interactive visualisations, and flexible download options, enabling researchers to access and apply these data with minimal need for additional engineering or background reading.

### Repertoire diversity across datasets

To evaluate the diversity of the antibody repertoires, we performed sequence identity clustering on the heavy and light chain variable regions with respect to IMGT variable region alignment. The raw number of sequences and the resulting cluster counts for each dataset, chain type, and region are detailed in **Table 2**. This analysis focused on the complementarity-determining regions (**CDR1, CDR2, CDR3**) and the framework regions (**FW**) using sequence identity thresholds of 90%, 80%, and 70%. To compare the heterogeneity of antibody responses, we calculated a normalized diversity percentage (number of unique clusters / total sequences), which is plotted in **Figure 2**. The results, as visualized in **Figure 2**, consistently demonstrate that the **CDR3** region of the heavy chain is the most diverse component of the antibody variable domain across all datasets and identity thresholds. For instance, in the top panel of **Figure 2** (90% identity threshold), 8 out of 11 datasets exhibit prominent heavy chain CDR3 diversity, with percentages exceeding 50%, while the light chain regions, by comparison, reflect their more limited contribution to general immune response variability by largely remaining below a 40% diversity threshold across the datasets. In contrast, the FW regions consistently show the lowest diversity (**Figure 2)**, which is expected given their conserved structural role. Furthermore, the data presented in **Table 2** and visualized in **Figure 2** reveal a wide spectrum of diversity profiles among the datasets, reflecting the nature of the specific immune response. The effect of the clustering strictness is also evident when comparing the panels of **Figure 2** from top to bottom; as the identity threshold is relaxed from 90% to 70%, the calculated diversity decreases across all regions due to the merging of sequences into fewer, more heterogeneous clusters. This comparative analysis highlights the varied complexity of antibody repertoires and confirms that our collection encompasses a broad range of immune responses suitable for robust immunological investigation.

**Table 2.** Sequence and Cluster Counts for Antibody Repertoires. This table provides a detailed breakdown of the total number of sequences (records) and the corresponding number of unique sequence clusters for each IMGT region (cdr1, cdr2, cdr3, fw). Data are presented for both heavy and light chains across all datasets, clustered at sequence identity thresholds of 70%, 80% and 90%.

|  | IMGT region |  | cdr1 |  |  | cdr2 |  |  | cdr3 |  |  | fw |  |  |
| --- | --- | --- | --- | --- | --- | --- | --- | --- | --- | --- | --- | --- | --- | --- |
|  | Identity threshold |  | 70% | 80% | 90% | 70% | 80% | 90% | 70% | 80% | 90% | 70% | 80% | 90% |
| Dataset | chain type | records |  |  |  |  |  |  |  |  |  |  |  |  |
| Acinetobacter baumannii #1 | heavy | 6144 | 326 | 529 | 782 | 180 | 305 | 490 | 2647 | 3613 | 4271 | 179 | 410 | 936 |
|  | light | 4711 | 251 | 298 | 556 | 154 | 155 | 155 | 664 | 1028 | 1531 | 196 | 453 | 1234 |
| Malaria CSP #1 | heavy | 16643 | 1031 | 1824 | 2943 | 872 | 1515 | 2583 | 6097 | 8400 | 9970 | 425 | 1112 | 3115 |
|  | light | 16425 | 899 | 1180 | 2072 | 606 | 629 | 658 | 1517 | 2501 | 3905 | 489 | 1197 | 3473 |
| Malaria CSP #2 | heavy | 21429 | 996 | 1738 | 2840 | 878 | 1535 | 2592 | 6498 | 9073 | 11251 | 384 | 919 | 2668 |
|  | light | 19845 | 903 | 1165 | 2032 | 618 | 645 | 684 | 1557 | 2480 | 3779 | 475 | 1083 | 2975 |
| Malaria CSP #3 | heavy | 5884 | 467 | 843 | 1381 | 456 | 789 | 1284 | 2228 | 2855 | 3306 | 188 | 480 | 1281 |
|  | light | 5997 | 530 | 708 | 1226 | 371 | 389 | 417 | 758 | 1234 | 1890 | 224 | 507 | 1413 |
| Pertussis | heavy | 11290 | 1215 | 2203 | 3517 | 986 | 1760 | 3003 | 3165 | 4033 | 4798 | 367 | 1031 | 3080 |
|  | light | 11290 | 843 | 1156 | 2072 | 561 | 582 | 621 | 1134 | 1906 | 2983 | 278 | 748 | 2383 |
| RSV #1 | heavy | 10368 | 589 | 1009 | 1558 | 432 | 747 | 1240 | 4315 | 6120 | 7295 | 182 | 507 | 1469 |
|  | light | 9697 | 548 | 751 | 1271 | 390 | 419 | 460 | 846 | 1327 | 2044 | 250 | 606 | 1730 |
| RSV #2 | heavy | 1051 | 134 | 229 | 352 | 130 | 205 | 308 | 509 | 595 | 656 | 39 | 90 | 236 |
|  | light | 1011 | 129 | 165 | 263 | 124 | 129 | 134 | 242 | 339 | 467 | 59 | 117 | 262 |
| SARS-CoV-2 #1 | heavy | 20189 | 1348 | 2422 | 3738 | 986 | 1699 | 2858 | 4500 | 6351 | 8080 | 557 | 1445 | 3728 |
|  | light | 25970 | 1048 | 1465 | 2762 | 431 | 437 | 440 | 1788 | 3177 | 5305 | 431 | 1188 | 3998 |
| SARS-CoV-2 #2 | heavy | 6659 | 591 | 977 | 1417 | 314 | 529 | 875 | 810 | 1040 | 1359 | 228 | 563 | 1315 |
|  | light | 9830 | 492 | 649 | 1224 | 215 | 217 | 217 | 621 | 1091 | 1766 | 180 | 524 | 1688 |
| Influenza #1 | heavy | 3949 | 493 | 807 | 1159 | 261 | 453 | 717 | 1744 | 2184 | 2409 | 120 | 345 | 975 |
|  | light | 3852 | 361 | 475 | 786 | 239 | 247 | 259 | 481 | 790 | 1189 | 161 | 380 | 1153 |
| Typhi #1 | heavy | 4435 | 344 | 639 | 1063 | 416 | 717 | 1204 | 1959 | 2440 | 2772 | 90 | 195 | 709 |
|  | light | 5005 | 345 | 447 | 796 | 269 | 278 | 287 | 674 | 1124 | 1706 | 100 | 186 | 543 |

**Figure 2.**
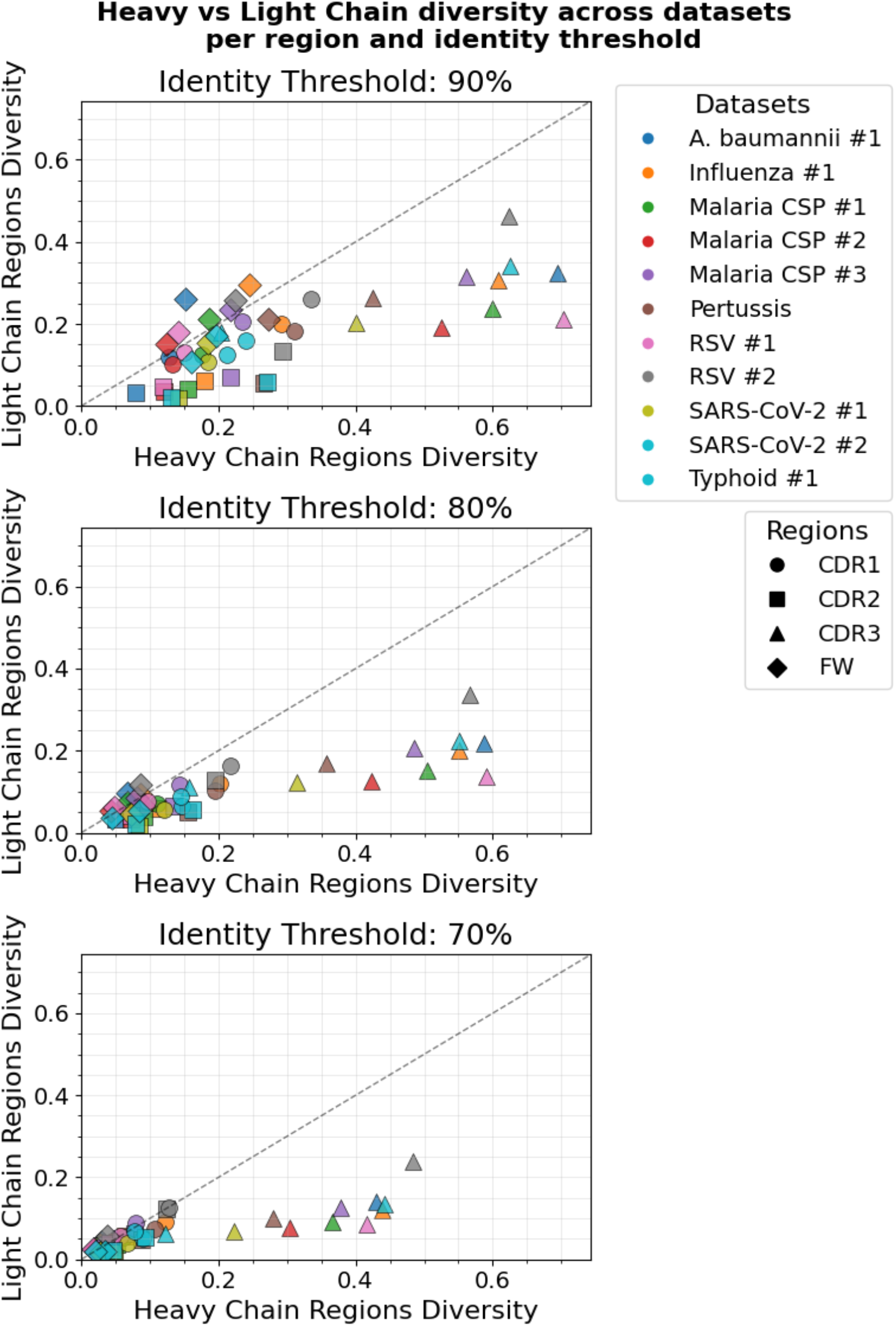
Comparative Diversity of Antibody Regions Across Datasets. The diversity calculated as the ratio of unique clusters to total chain sequences is shown for complementarity-determining regions (CDR1, CDR2, CDR3) and framework regions (FW). The analysis is presented for **11** different antigen-specific datasets at three distinct sequence identity thresholds: **90%** (top), **80%** (middle), and **70%** (bottom). The results highlight the consistently higher diversity of the CDR3 region and the variable diversity profiles across the different datasets.

## Discussion

Despite the rapid growth of public antibody databases, there is currently no centralized resource that captures antibody discovery data to such a large scale. Most existing public resources of therapeutic antibody primarily curate clinically successful antibodies from sparse origins, representing only a small fraction of the total discovery funnel and seldom include negative or low-binding clones^7 21^.

KyDab addresses this gap by providing a comprehensive antibody discovery dataset generated under high quality standard and consistent workflows. All data were produced using a well-established humanized mouse platform with standardized immunisation, single-cell sequencing, selection and expression pipelines across multiple studies. This level of upstream consistency is rarely achieved in public antibody resources. Importantly, the dataset reflects the full process of antibody down-selection, from large repertoires of paired VH–VL sequences to a defined high-potential subset advanced to experimental testing. Such data are particularly valuable for AI applications aimed at improving the success rate of experimental validation. In addition, the antigen diversity in KyDab also provides a more challenging and realistic benchmark for evaluating model robustness across targets.

As with all antibody discovery workflows, the dataset reflects biases introduced by bioinformatics-guided down-selection. Sequences advanced to expression are enriched for expanded lineages, higher somatic mutation levels and predicted developability, leading to an overrepresentation of true binders and high-affinity antibodies. Nevertheless, KyDab includes both positive and negative experimental outcomes. The availability of negative data is critical for proper model calibration and for evaluating false-positive behavior in the presence of selection bias.

The long-term goal of AI-driven antibody discovery is to improve success rate, reduce cost and development time by shifting key decision-making steps from experimental screening to computational prediction. While AI has demonstrated impressive progress in antibody in silico generation, its application to virtual screening and affinity prediction remains constrained by the lack of large, high-quality antibody discovery datasets^3^. By releasing KyDab as an open resource, we aim not only to support this next phase of AI-empowered antibody discovery, but also to demonstrate the value of sharing structured, full-funnel discovery data. We hope this effort will encourage broader data contributions from across the antibody discovery community, particularly from industrial organisations, to enable the development of more robust and generalisable AI models through collective progress.

## Acknowledgments

The authors thank Donald Drake and Matthew Massett for manuscript review; Aaron Hanley and Josefin Bartholdson-Scott for assistance with dataset preparation; Janet Tse and Juliette Goutorbe for preparing the Terms and Conditions; Jemima Paterson, Andrew Phillips, Danai Koftori and Jean-Philippe Mametz for support with coordination and digital advice.

